# Postsynaptic serine racemase regulates NMDA receptor function

**DOI:** 10.1101/2020.06.16.155572

**Authors:** Jonathan M. Wong, Oluwarotimi Folorunso, Eden V. Barragan, Cristina Berciu, Theresa L. Harvey, Michael R. DeChellis-Marks, Jill R. Glausier, Matthew L. MacDonald, Joseph T. Coyle, Darrick T. Balu, John A. Gray

## Abstract

D-serine is the primary NMDA receptor (NMDAR) co-agonist at mature forebrain synapses and is synthesized by the enzyme serine racemase (SR). However, our understanding of the mechanisms regulating the availability of synaptic D-serine remains limited. Though early studies suggested D-serine is synthesized and released from astrocytes, more recent studies have demonstrated a predominantly neuronal localization of SR. More specifically, recent work intriguingly suggests that SR may be found at the postsynaptic density, yet the functional implications of postsynaptic SR on synaptic transmission are not yet known. Here, we show an age-dependent dendritic and postsynaptic localization of SR and D-serine by immunohistochemistry and electron microscopy in mouse CA1 pyramidal neurons, as well as the presence of SR in human hippocampal synaptosomes. In addition, using a single-neuron genetic approach in SR conditional knockout mice, we demonstrate a cell-autonomous role for SR in regulating synaptic NMDAR function at Schaffer collateral (CA3)-CA1 synapses. Importantly, single-neuron genetic deletion of SR resulted in the elimination of LTP at one month of age. Interestingly, there was a restoration of LTP by two months of age that was associated with an upregulation of synaptic GluN2B. Our findings support a cell-autonomous role for postsynaptic neuronal SR in regulating synaptic NMDAR function and suggests a possible autocrine mode of D-serine action.

## Introduction

NMDA receptors (NMDARs) are glutamate receptors, which have a property unique among ion channels: they require a co-agonist for channel opening (Johnson and Ascher, 1987; Kleckner and Dingledine, 1988). In addition to glutamate binding to the GluN2 subunits, either glycine or D-serine must bind to the GluN1 subunits. D-serine is the primary co-agonist at most mature forebrain synapses, including the Schaffer collateral-CA1 synapse in the hippocampus (Mothet et al., 2000; Papouin et al., 2012; Le Bail et al., 2015) and is synthesized in the brain by the enzyme serine racemase (SR), which converts L-serine to D-serine (Wolosker et al., 1999). However, our understanding of the mechanisms regulating the availability of synaptic D-serine remains limited.

Early studies suggested that D-serine is synthesized and released by astrocytes leading to the labeling of D-serine as a “gliotransmitter” (Schell et al., 1995; Schell et al., 1997; Wolosker et al., 1999; Panatier et al., 2006). However, more recent studies have demonstrated a predominantly neuronal localization (Wolosker et al., 2016). Indeed, germline SR knock-out (KO) mice have been used to validate SR antibody specificity, identifying a preferential expression of SR in neurons in rodent and human brains (Kartvelishvily et al., 2006; Miya et al., 2008; Basu et al., 2009; Ding et al., 2011; Balu et al., 2014; Balu et al., 2018). This neuronal localization of SR was further supported by *in situ* hybridization (Yoshikawa et al., 2007) and in transgenic mice where the SR coding region was replaced by GFP (Ehmsen et al., 2013). Convincingly, genetic deletion of SR from pyramidal neurons, but not from astrocytes, leads to reduction of brain D-serine concentration (Benneyworth et al., 2012; Ishiwata et al., 2015), and impairment of long term potentiation (LTP) at CA3-CA1 synapses (Benneyworth et al., 2012; Perez et al., 2017). Furthermore, biochemical evidence from adult rat brain demonstrated the presence of SR in synaptosomes (Balan et al., 2009), and SR has been found to co-localize and co-immunoprecipitate with postsynaptic density protein 95 (PSD-95) in cortical neuronal cultures, supporting a postsynaptic localization of SR (Ma et al., 2014; Lin et al., 2016). However, the functional implications of postsynaptic SR on synaptic transmission are not known.

Here, we demonstrate dendritic and postsynaptic localization of SR and D-serine by immunohistochemistry and electron microscopy in hippocampal CA1 neurons. In addition, using a single-neuron genetic approach in SR conditional knockout mice, we demonstrate a cell-autonomous role for SR in regulating synaptic NMDAR function. Importantly, single-neuron genetic deletion of SR resulted in the elimination of CA3-CA1 LTP at one month of age. Interestingly, LTP was restored by two months with a concomitant upregulation of synaptic GluN2B. This evidence supports a cell-autonomous role for postsynaptic neuronal SR in regulating synaptic function and suggests a possible autocrine mode of D-serine action.

## Materials and Methods

### Animals

The floxed (fl) SR construct was generated as previously described (Basu et al., 2009; Benneyworth et al., 2012). In this construct, the first coding exon (exon 3) is flanked by loxP sites, which results in excision of the intervening sequence upon exposure to Cre recombinase. These *Srr*^fl/fl^ mice are maintained on a C57B/L6J background. Mice were grouped housed in polycarbonate cages and maintained on a 12-hour light/dark cycle. Animals were given access to food and water *ad libitum*. The University of California Davis and McLean Hospital Institutional Animal Care and Use Committee approved all animal procedures.

### Immunofluorescence

Postnatal day 16 (P16) and two month-old wild-type C57B/L6 mice were deeply anesthetized, briefly intracardially perfused with cold PBS (0.5M phosphate buffer, NaCl, pH 7.4), followed by 4% paraformaldehyde (Electron Microscopy Sciences, Cat # 19202) and then cryopreserved in 30% sucrose/PBS at 4°C. Brains were sectioned at 30 μm using a Leica SM 2010R Microtome or Microme HM 505E Cryostat and stored in a cryoprotectant solution (ethylene glycol, glycerol, 0.5M PB, NaCl, KCl, in dH_2_O) at −20°C. Free floating sections were washed three times with PBS, then pre-incubated with permeabilizing agent 0.3% Triton X in PBS for 30 min. Sections were incubated in blocking buffer (20% donkey serum, 1% BSA, 0.1% glycine, 0.1% lysine in PBS) for 1hr. This was followed by overnight incubation at 4°C with primary antibodies in incubation buffer (5% donkey serum, 1% BSA, 0.1% glycine, and 0.1% lysine). Primary antibodies used in this study were: mouse anti-serine racemase (BD biosciences, cat #612052), rabbit anti-PSD95 (Abcam, cat #12093), rabbit anti-MAP2 (Abcam, cat# 5622). Sections were washed three times with PBS, then incubated with the appropriate secondary antibodies in incubation buffer for 1hr. The following isotype-specific secondary antibodies were used: donkey anti-mouse Alexa 488, donkey anti-rabbit Alexa 568 with, donkey anti-rabbit Alexa 647, goat anti-mouse Alexa 488, and goat anti-rabbit Alexa 568. After secondary antibody incubation, tissues were washed three times with 1x PBS. Before mounting on Fisherbrand Superfrost Plus glass microscope slides (Fisher Scientific), free-floating slices were rinsed with water and counterstained using the nuclear maker DAPI. Finally, glass slides were covered using cover glass with ProLong Gold anti-fade (Invitrogen). Confocal images were acquired using a Leica SP8 confocal microscope (20x/40x objectives) and Z-series stack confocal images were taken at fixed intervals using consistent settings.

### Electron Microscopy

Two month-old mice were deeply anesthetized, briefly intracardially perfused with cold PBS (0.5M phosphate buffer, NaCl, pH 7.4), followed by either (3% glutaraldehyde, 1% paraformaldehyde and 0.2% sodium metabisulfite in 0.1M phosphate buffer, pH7.4) for D-serine staining (Balu et al., 2014) or (4% paraformaldehyde and 0.5% gluteraldehyde in 0.1M PB pH7.4) for SR staining. Brains were post-fixed in either CaCl_2_, 0.1% sucrose, 3% glutaraldehyde, 1% PFA in dH_2_O (pH 7.4; D-serine staining) or 4% PFA, 0.1% sucrose, 0.5% glutaraldehyde (pH 7.4; SR staining). Brains were sectioned using a Leica VT1200S Vibratome at 40 μm.

#### Nanogold staining

Hippocampal sections were post fixed in 1% OsO_4_, dehydrated in graded ethanol series and extra dry acetone, and flat embedded in Embed 812 resin. Ultrathin sections (~80 nm) were collected on Formvar-coated slot grids. Sections were quickly imaged to determine proper orientation of sections. Sections were washed with filtered (0.05M TBS pH 7.4 with 0.15% glycine) for 10 mins, and then incubated with filtered (2% normal goat serum in 0.05M TBS, pH 7.4) for 1 hour to block non-specific protein binding sites. Sections were washed again (0.05M TBS pH 7.4) and incubated with rabbit anti-D-serine antibody (1:1000, Abcam, #6472) with 0.05mM L-serine-BSA-glutaraldehyde conjugate overnight at RT, adjusted from (Balu et al., 2014). We previously validated this D-serine IHC protocol using SRKO tissue to demonstrate the necessity of L-serine blocking conjugate inclusion to prevent antibody cross reactivity with L-serine, which is highly expressed in astrocytes (Yang et al., 2010). Following incubation, sections were washed (0.05M TBS pH 7.4) and incubated with goat anti-rabbit IgG H&L (10nm Gold; 1:20, Abcam cat# 39601) in filtered 2% normal goat serum (0.05 M TBS; pH 8.2). Before being imaged, sections were post-stained with uranyl acetate (saturated solution) and Reynold’s lead citrate.

#### DAB (3,3’-Diaminobenzidine)

Endogenous peroxidases were quenched by incubating sections with 0.3% H_2_O_2_ in 0.01M PBS for 15mins. Sections were washed in wash buffer 1 (0.01% Triton X-100 in 0.01M PBS) three times and then incubated with 0.05% fresh NaBH_4_ containing 0.1% glycine for 30mins to quench aldehydes. Sections were washed with 0.01M PBS three times, blocked in (4% BSA, 10% normal goat serum, 0.01% triton-X-100 in 0.01M PBS) for one hour at RT. Sections were rinsed once with wash buffer 1 and incubated for two days at 4°C with mouse anti-SR antibody (1:1,000; BD Biosciences; cat #612052). Sections were washed with wash buffer 2 (2% BSA, 2% normal goat serum, 0.01% triton-X-100 in 0.01M PBS). Goat anti-mouse IgG-biotinylated was used to incubate sections for 2 hours at RT. Sections were washed with wash buffer 2. Sections were incubated for 2 hours at RT with strepavidin–HRP (Invitrogen; Cat #434323). Sections were washed twice in 0.1M phosphate buffer, incubated sections with DAB - H_2_O_2_ (Vector; Cat #4105) made in 0.01M Na cacodylate, and then washed twice with 0.1M PB.

#### Imaging

Sections were imaged on a JEOL JEM-1200 EX II with an 1k CCD camera, and on a Tecnai F20 (200 keV) transmission electron microscope (FEI, Hillsboro, OR) and recorded using a 2K × 2K charged-coupled device (CCD) camera, at 17,500X magnification (1.12 nm pixel size). For large overviews, we acquired montages of overlapping high-magnification images in an automated fashion using the microscope control software SerialEM (Mastronarde, 2005).

### Proteomics

Hippocampal grey matter (CA1 and DG including pyramidal and granule cell layers and neuropil, 20 mg) was dissected from brain specimens of three postmortem human subjects with no evidence of lifetime psychiatric or neurologic disorders (Glausier et al., 2020). Tissue homogenates and synaptosomes were prepared using the Syn-PER™ Synaptic Protein Extraction Reagent (Thermo) per manufacture instructions. 60 μg total protein (as measured by Micro BCA™ Protein Assay, Thermo) from each sample were digested with trypsin on S-Traps Minis (Protifi) following manufacture protocol. 10 μg of digested peptides from each sample were labeling using TMT10plex™ Isobaric Mass Tagging Kit (Thermo) at a 4:1 (wt:wt) ratio of label-to-peptide [based on (Zecha et al., 2019)]. The labeled-peptides were then pooled and fractioned using the Pierce™ High pH Reverse-Phase Peptide Fractionation Kit (Thermo), per modified manufacture instructions. In total, nine fractions were eluted with the following acetonitrile-triethylamine gradients: (1) 5.0%, (2) 10.0%, (3) 12.5%, (4) 15.0%, (5) 17.5%, (6) 20.0%, (7) 22.5%, (8) 25.0%, and (9) 50%. Approximately 1 μg of peptides from each fraction were resolved on an EASY C18 column (1.7μm, 2.1×50cm) at 300 nL/min with an UltiMate™ 3000 RSLCnano HPLC system over a 90-min gradient and analyzed on Orbitrap Eclipse™ Tribrid™ MS operated in MS^3^ with real-time search and synchronous precursor selection. Peptide/protein identification and quantification was performed in Proteome Discoverer ™ (v2.4, Thermo).

### Electrophysiology

#### Postnatal viral injection

Neonatal [P0-P1] *Srr*^fl/fl^ mice of both sexes were stereotaxically injected with high-titer rAAV1-Cre:GFP viral stock (~1-5 x 10^12^ vg/mL) with coordinates targeting CA1 of hippocampus as previously described (Gray et al., 2011). Transduced neurons were identified by nuclear GFP expression. Cre expression was generally limited to the hippocampus within a sparse population of CA1 pyramidal neurons.

#### Acute slice preparation

Mice older than P30 were anesthetized with isoflurane and transcardially perfused with ice-cold artificial cerebrospinal fluid (ACSF), containing (in mM) 119 NaCl, 26.2 NaHCO_3_, 11 glucose, 2.5 KCl, 1 NaH_2_PO_4_, 2.5 CaCl_2_ and 1.3 MgSO_4_. Modified transverse 300 μm slices of dorsal hippocampus were prepared by performing a ~10^0^ angle blocking cut of the dorsal portion of each cerebral hemisphere (Bischofberger et al., 2006) then mounting the cut side down on a Leica VT1200 vibratome in ice-cold cutting buffer. Slices were incubated in 32°C NMDG solution containing (in mM) 93 NMDG, 93 HCl, 2.5 KCl, 1.2 NaH_2_PO_4_, 30 NaHCO_3_, 20 HEPES, 25 glucose, 5 sodium ascorbate, 2 thiourea, 3 sodium pyruvate, 10 MgSO_4_, and 0.5 CaCl_2_ (Ting et al., 2018) for 15 mins, transferred to room temperature ACSF, and held for at least 1 hr before recording. Mice younger than P30 were anesthetized in isoflurane and decapitated. Brains were rapidly removed and placed in ice-cold sucrose cutting buffer, containing the following (in mM): 210 sucrose, 25 NaHCO_3_, 2.5 KCl, 1.25 NaH_2_PO_4_, 7 glucose, 7 MgCl_2_, and 0.5 CaCl_2_. Slices were cut as described above. Slices were recovered in 32°C ACSF. All solutions were vigorously perfused with 95% O_2_ and 5% CO_2_. Slices were transferred to a submersion chamber on an upright Olympus microscope, perfused in room temperature ACSF containing picrotoxin (0.1 mM), and saturated with 95% O_2_ and 5% CO_2_. CA1 neurons were visualized by infrared differential interference contrast microscopy, and GFP+ cells were identified by epifluorescence microscopy.

#### Whole-cell patch clamp

Cells were patched with 3-5 MΩ borosilicate pipettes filled with intracellular solution containing (in mM) 135 cesium methanesulfonate, 8 NaCl, 10 HEPES, 0.3 Na-GTP, 4 Mg-ATP, 0.3 EGTA, and 5 QX-314 (Sigma, St Louis, MO). Series resistance was monitored and not compensated, and cells were discarded if series resistance varied more than 25%. All recordings were obtained with a Multiclamp 700B amplifier (Molecular Devices, Sunnyvale, CA), filtered at 2 kHz, and digitized at 10 Hz. All EPSCs were evoked at 0.1Hz. AMPA receptor EPSCs were recorded at −70mV. NMDAR-EPSCs were recorded in the presence of 10μM NBQX at +40mV with the exception of d-serine wash experiments, which were recorded at −40mV (Bergeron et al., 1998; Basu et al., 2009). Dual-whole cell recordings are the average response of approximately 40 sweeps, the typical recording length. Paired pulse response (PPR) was performed with two sequential EPSCs at a 50 ms interval. LTP was induced by depolarization to 0mV paired with 1Hz stimulation for 90 s. Ro25-6981 (Ro25) and d-serine wash experiments recorded NMDAR-EPSCs by coming to a steady 5 min baseline before proceeding with wash. EPSC amplitude was determined by measuring the peak of the response compared to pre-stimulation baseline. All summary amplitude graphs after a time course experiment average the last 5 min of data compared to baseline. EPSC charge transfer was determined by measuring the area of the response compared to pre-stimulation baseline. EPSC decay time was measured as the time between EPSC peak amplitude and 63% decay from the peak. Analysis was performed with the Clampex software suite (Molecular Devices).

### Experimental design and statistical analysis

All data represents the mean ± SEM of n = number of neurons or pairs of neurons. In electrophysiology experiments, two to three data points were typically acquired per mouse. Experiments include both males and females. Data were analyzed using Clampfit 10.4 (Axon Instruments) and Prism 8 software (GraphPad). Paired amplitude data were analyzed with paired two-tailed *t* test and unpaired two-tailed *t* test where p<0.05 was considered significant. For TEM, micrographs were visualized on IMOD software (Kremer et al., 1996), and clearly defined synapses (pre- and post-neurons) were identified and all postsynaptic neurons were manually counted for nanogold labeled D-serine. No D-serine was detected in the presynaptic neuron. For SR quantification, in Image J, a region of interest (ROI) was drawn around fifty synapses for both SR labeled and control (no antibody) sections; pixel intensity was quantified per ROI for each synapse.

## Results

### Postsynaptic localization of serine racemase and D-serine

Biochemical evidence from adult rat brain demonstrated the presence of SR in synaptic compartments (Balan et al., 2009), while in primary mouse neuronal cortical cultures, SR and D-serine co-localize with PSD-95 and NMDARs (GluN1) in postsynaptic glutamatergic synapses, but not in pre-synaptic terminals (Ma et al., 2014; Lin et al., 2016). Furthermore, SR expression is developmentally regulated in the mouse brain, with low levels early postnatally and peaking ~P28 (Miya et al., 2008; Basu et al., 2009). In the CA1 subfield of dorsal hippocampus at P16 (Fig 1A,C), SR was not detected in either the pyramidal cell layer or apical dendrites of *stratum radiatum* (*s.r*.) and *lanconosum moleculare* (*l.m*.). By two months of age, there is robust SR expression throughout CA1, including high levels in apical dendrites as demonstrated by co-localization with microtubule-associated protein 2 (MAP2; Fig 1B,D). Using immuno-electron microscopy (EM), we show that SR is present in CA1 apical dendrites of *s.r*., with no SR immunoreactivity in control sections, in which only the primary SR antibody was omitted (Fig 1E,F).

**Figure 1:**
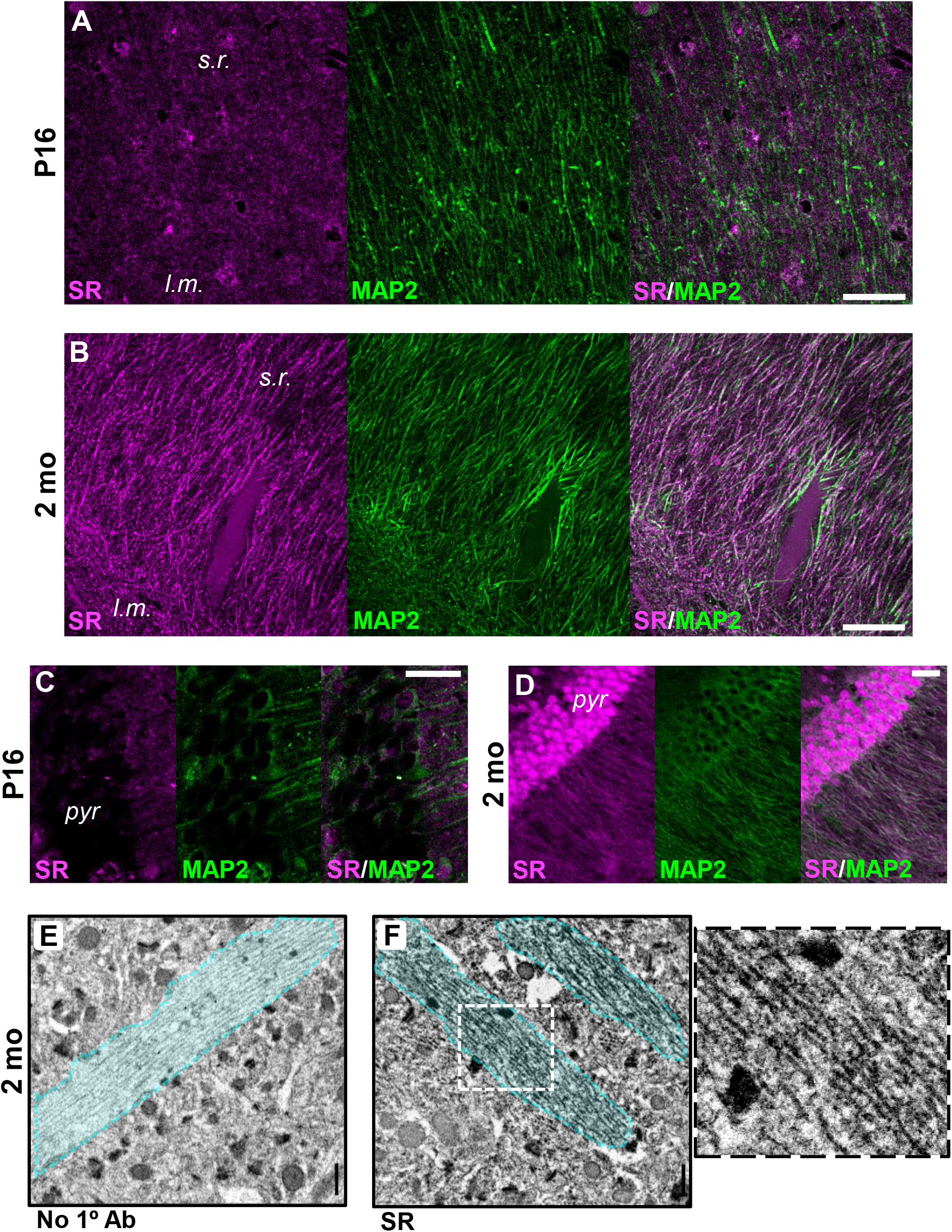
SR is present in the apical dendrites in CA1. **(A-D)** Representative confocal images showing co-localization of SR immunofluorescence (magenta) with microtubule associated protein 2 (MAP2; green) in the apical dendrites of CA1 pyramidal neurons in *stratum radiatum* (*s.r*.; A,B) and *stratum pyramidale* (*pyr*; C,D) at either P16 (A,C) or 2 months (B,D); *lacunosum moleculare* (*l.m*.). Scale bars 10μM. **(E-F)** TEM micrographs showing SR DAB photoreaction product in CA1 apical dendrites at 2 months (n = 3 mice). Scale bars = 500nM

We also examined whether SR was localized to postsynaptic compartments in *s.r*. of CA1. We found more intense SR immunoreactivity at postsynaptic densities (PSDs) in SR-antibody stained sections compared to control sections with no primary antibody, demonstrating the presence of SR in dendritic spines (Fig 2A-C; No 1° Ab, 78.0 ± 11.4 (SD), n = 48; SR Ab, 110.6 ± 15.3 (SD), n = 50). However, we did not detect SR immunoreactivity in presynaptic compartments (Fig 2B). Using dual-antigen immunofluorescence, we also demonstrate co-localization of SR with the PSD marker, PSD-95 in CA1 (Fig 2D). Finally, mass spectrometry confirmed the presence of SR in synaptosomes (MacDonald et al., 2019) prepared from human hippocampus, along with other well-known pre- and post-synaptic markers (Fig 2E, Extended Data Table 2-1).

**Figure 2:**
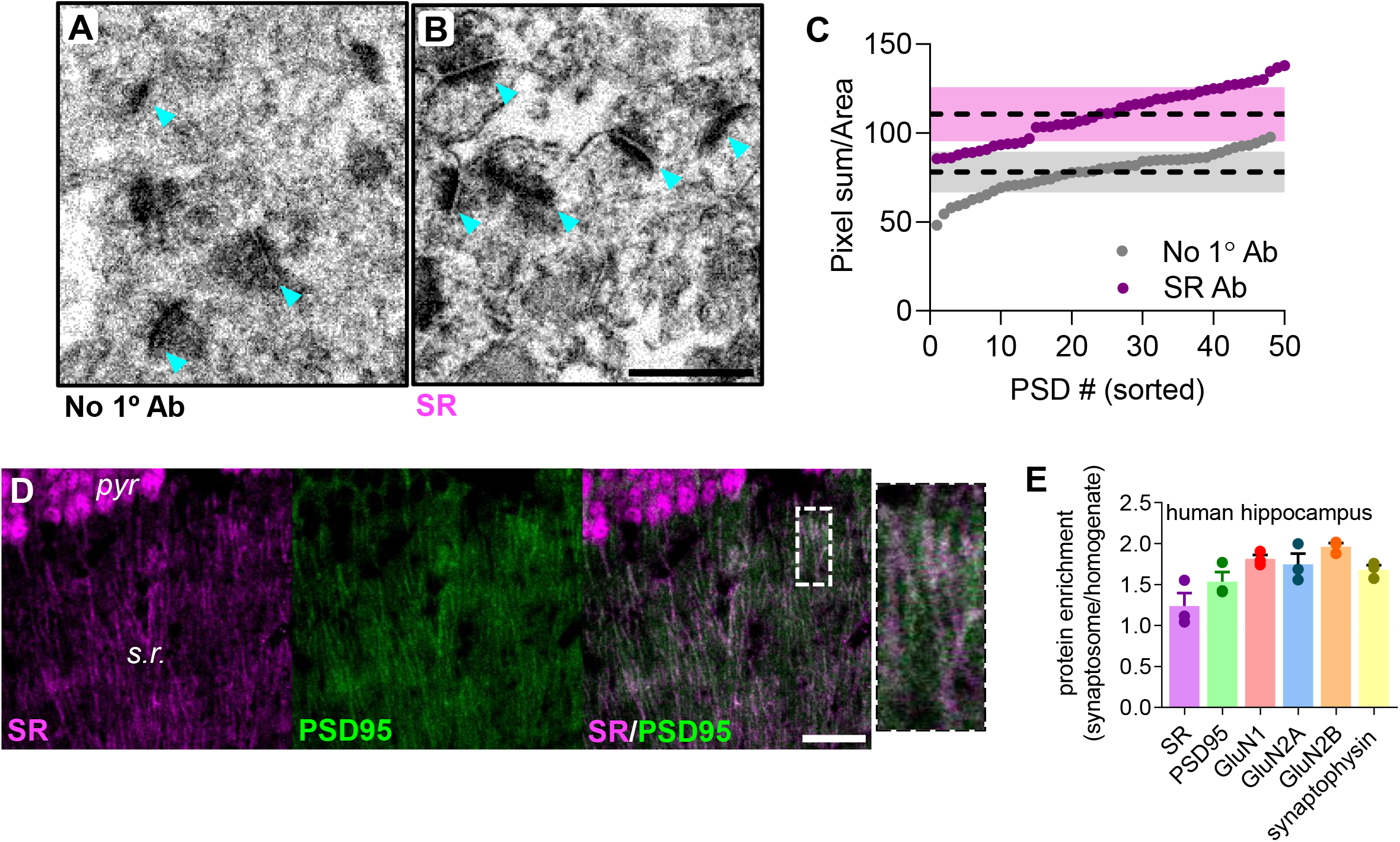
SR is enriched in the PSD in CA1. **(A,B)** TEM micrographs showing SR DAB photoreaction product in hippocampal synapses of CA1 stratum radiatum. Scale bars = 500nM; n = 1 mouse. **(C)** Intensity of DAB product at individual postsynaptic densities (PSD) as pixel intensity per region of interest (ROI), sorted to show range. Dashed lines represent average pixel intensity ± SD in shaded bars (No 1° Ab, 78.0 ± 11.4, n = 48; SR Ab, 110.6 ± 15.3, n = 50). **(D)** Representative confocal images of SR immunofluorescence (magneta) showing colocalization with PSD95 (green) at 2 months. Scale bars 10μM; n = 3 mice; *stratum pyramidale* (*pyr*); *stratum radiatum* (*s.r*.). **(E)** Mass spectrometry identifying enriched proteins in human hippocampal synaptosomes (n = 3 subjects).

Since we detected the enzyme SR in dendrites and spines, we next examined whether the NMDAR co-agonist D-serine is also localized to these compartments. Using immuno-EM and a D-serine immunostaining protocol that we previously validated using SRKO mice (Balu et al., 2014), we observed high numbers of nanogold particles in CA1 *s.r*. dendrites, while we did not detect any dendritic nanogold particle binding in control sections that omitted only D-serine primary antibody (Fig 3A-C). Finally, we detected nanogold particles post-synaptically in dendritic spines in CA1, but not in pre-synaptic compartments (data not shown) or in control sections that were not incubated with the D-serine primary antibody (Fig 3D-F).

**Figure 3:**
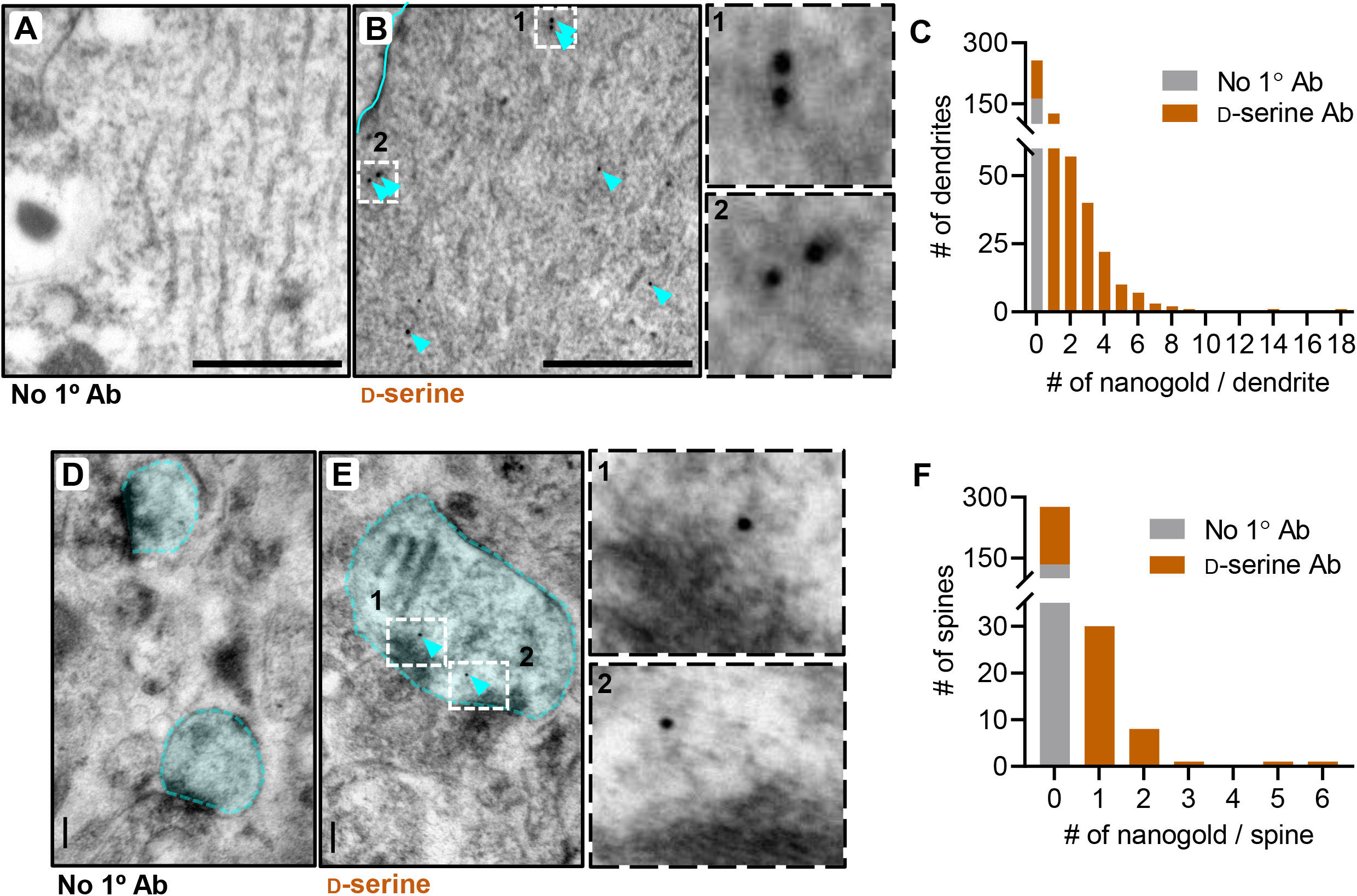
D-serine is present in dendrites and spines in CA1. **(A,B)** TEM micrographs showing D-serine nanogold particles in hippocampal CA1 apical dendrites at 2 months. **(C)** Histogram showing the number of D-serine nanogold particles in dendritic segments (n = 1 mouse; 150-500 dendrites). **(D,E)** TEM micrographs showing D-serine nanogold particles in dendritic spines in CA1 *stratum radiatum* at 2 months. **(F)** Histogram showing the number of D-serine nanogold particles in dendritic spines (n = 1 mouse; 130-300 dendritic spines). Scale bar - 100nM

### Single-neuron genetic deletion of serine racemase does not alter synaptic function in P14-21 CA1

To examine the physiological function of postsynaptic SR, we utilized a single-neuron genetic approach in SR conditional knockout mice. Here, SR was removed in a sparse subset of CA1 pyramidal neurons by postnatal day 0 (P0) stereotaxic injection of adeno-associated virus, serotype 1 expressing a Cre recombinase GFP fusion protein (AAV1-Cre:GFP) into floxed SR (*Srr*^fl/fl^) mice (Fig 4A). This mosaic transduction allows for simultaneous whole-cell recordings from Cre-expressing (Cre+) and neighboring untransduced neurons (control) (Fig 4B), providing a rigorous comparison of the cell-autonomous effects of SR deletion while controlling for presynaptic input (Gray et al., 2011).

**Figure 4:**
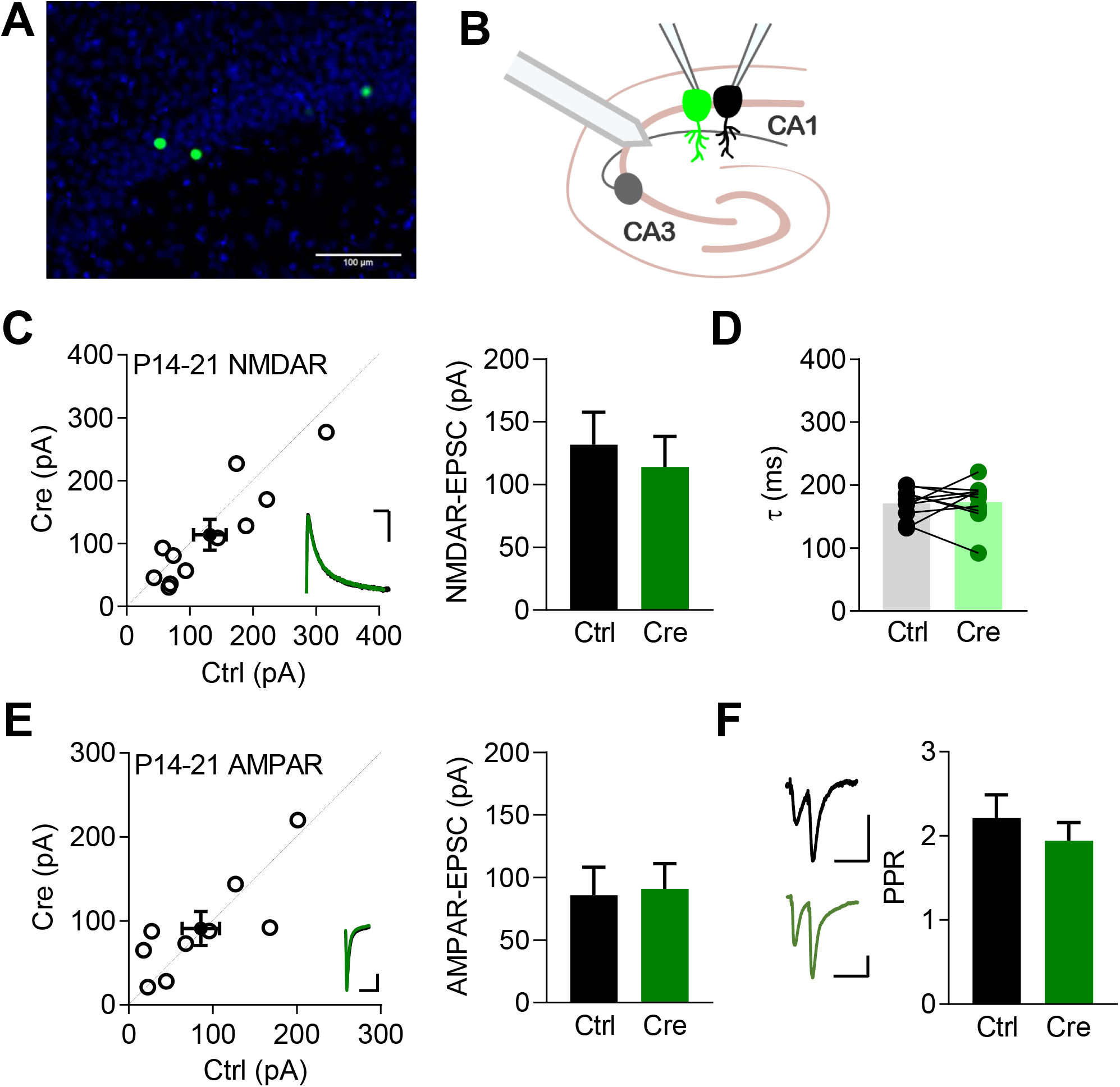
No effects at P14-21 following single-neuron SR deletion. **(A)** Representative image of the sparse transduction of CA1 pyramidal cells by AAV1-Cre:GFP counterstained by DAPI. Scale bar indicates 100μm. **(B)** Schematic of experimental setup. Dual whole-cell EPSC recordings from Schaffer collateral stimulation were made from neighboring transduced and control CA1 pyramidal cells. **(C)** Scatterplot of paired neuronal recordings of P14-21 NMDAR-EPSCs (open circles) and averaged pair ± SEM (solid circle). Sample trace bars indicate 200ms, 30pA. Average NMDAR-EPSC amplitudes for control (131.7 ± 25.9 pA, n = 11) and Cre:GFP+ neurons (114.0 ± 24.4 pA, n = 11; t(10) = 1.591, p = 0.1426, paired t test). **(D)** Average decay kinetics of NMDAR-EPSCs for control (170.0 ± 7.5 ms, n = 10) and Cre:GFP+ neurons (173.1 ± 10.8 ms, n = 10) from paired recordings in C (t(9) = 0.2197, p = 0.8310, paired t test). **(E)** Scatterplot of paired neuronal recordings of P14-21 AMPAR-EPSCs (open circles) and averaged pair ± SEM (solid circle). Sample trace bars indicate 200ms, 30pA. Average AMPAR-EPSC amplitudes for control (85.9 ± 22.3 pA; n = 9) and Cre:GFP+ neurons (91.0 ± 20.2 pA; n = 9; t(8) = 0.3917, p = 0.7055, paired t test). **(F)** Average paired pulse ratio for control (2.21 ± 0.28, n = 4) and Cre:GFP+ neurons (1.94 ± 0.21, n = 4; t(6) = 0.7694, p = 0.4708, t test). Sample trace bars indicate 100ms, 30pA.

Glycine is thought to be the primary synaptic NMDAR co-agonist at early developmental stages before being gradually supplanted by D-serine around the third postnatal week in CA1 (Le Bail et al., 2015). Therefore, we first assessed the contribution of postsynaptic SR to synaptic physiology at P14-21. In P14-21 mice, we found no difference in the NMDAR-EPSC amplitudes between control and Cre+ neurons (Fig 4C; control: 131.7 ± 25.9 pA, n = 11; Cre+: 114.0 ± 24.4 pA, n = 11; t(10) = 1.591, p = 0.1426, paired t test). Decay kinetics of NMDAR-EPSCs, measured as the time between the EPSC peak and 63% decay, were also unchanged (Fig 4D; control: 170.0 ± 7.5 ms, n = 10; Cre+: 173.1 ± 10.8 ms, n = 10; t(9) = 0.2197, p = 0.8310, paired t test). We also recorded AMPAR-EPSCs from P14-21 mice in a pairwise manner and likewise found no change in AMPAR-EPSCs (Fig 4E; control: 85.9 ± 22.3 pA, n = 9; Cre+: 91.0 ± 20.2 pA, n = 9; t(8) = 0.3917, p = 0.7055, paired t test). Finally, we recorded paired pulse ratios (PPR) with a 50 ms interval from control and Cre+ neurons and found no change in PPR (Fig 4F; 2.21 ± 0.28, n = 4; Cre+: 1.94 ± 0.21, n = 4; t(6) = 0.7694, p = 0.4708, t test). As expected, postsynaptic SR deletion has no observed effect on synaptic function in P14-21 CA1 consistent with glycine being the primary synaptic co-agonist at this timepoint.

### Single-neuron genetic deletion of serine racemase reduces NMDAR-EPSCs in P45-70 CA1

Next, we performed recordings around 2 months of age when D-serine is clearly the primary synaptic NMDAR co-agonist (Papouin et al., 2012; Le Bail et al., 2015). In P45-70 mice, we found that postsynaptic SR deletion decreased NMDAR-EPSCs in Cre+ neurons (Fig 5A; control: 144.1 ± 18.9 pA, n = 19; Cre+: 87.6 ± 11.7 pA, n = 19; t(18) = 3.365, p = 0.0034, paired t test). Importantly, AMPAR-EPSCs were unchanged indicating that the effect was specific to NMDARs (Fig 5B; control: 111.8 ± 19.0 pA, n = 14; Cre+: 85.4 ± 15.7 pA, n = 14; t(13) = 1.854, p = 0.0866, paired t test). PPR was also unchanged (Fig 5C; control: 1.83 ± 0.15, n = 9; Cre+: 1.70 ± 0.22, n = 9; t(16) = 0.4649, p = 0.6483, t test), suggesting that the NMDAR-EPSC decrease was postsynaptic in origin. Previous studies have demonstrated that acute removal of NMDAR co-agonists impairs synaptic plasticity (Papouin et al., 2012; Le Bail et al., 2015) and that germline and neuron-specific SR deletion reduces LTP (Basu et al., 2009; Benneyworth et al., 2012; Balu et al., 2013; Perez et al., 2017). Therefore, we assessed the effect of single cell postsynaptic deletion of SR on LTP. Surprisingly, we found that LTP was unchanged in Cre+ neurons (Fig 5D; control: 193.9 ± 22.1 %, n = 12; Cre+: 196.6 ± 15.1 %, n = 12; t(22) = 0.09961, p = 0.9216, t test).

**Figure 5:**
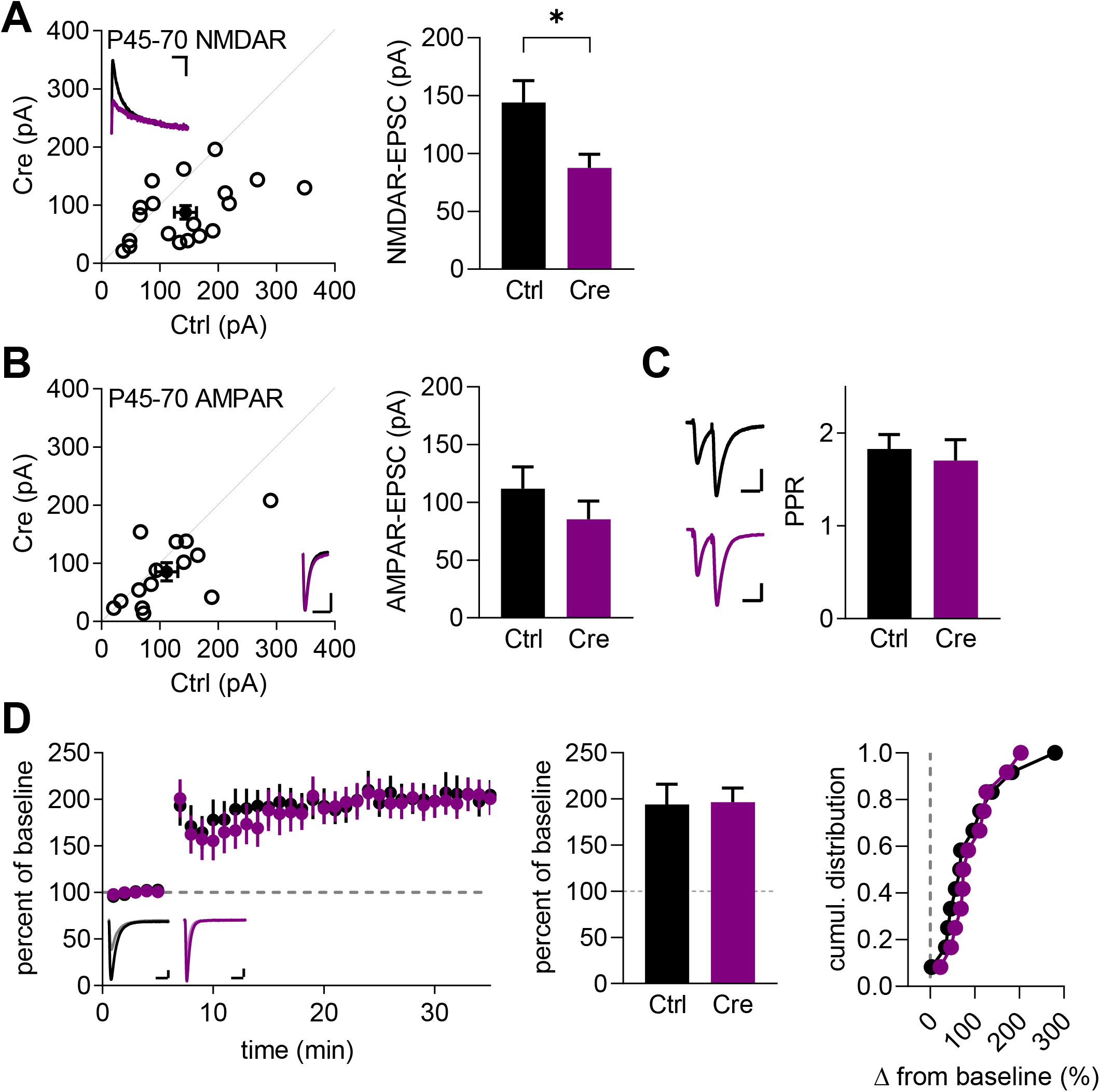
Reduced NMDAR-EPSCs at P45-70 following single-neuron SR deletion. **(A)** Scatterplot of paired neuronal recordings of P45-70 NMDAR-EPSCs (open circles) and averaged pair ± SEM (solid circle). Sample trace bars indicate 200ms, 30pA. Average NMDAR-EPSC amplitudes for control (144.1 ± 18.9 pA, n = 19) and Cre:GFP+ neurons (87.6 ± 11.7 pA, n = 19; t(18) = 3.365, p = 0.0034, paired t test). **(B)** Scatterplot of paired neuronal recordings of P45-70 AMPAR-EPSCs (open circles) and averaged pair ± SEM (solid circle). Sample trace bars indicate 200ms, 30pA. Average AMPAR-EPSC amplitudes for control (111.8 ± 19.0 pA, n = 14) and Cre:GFP+ neurons (85.4 ± 15.7 pA; t(13) = 1.854, p = 0.0866, paired t test). **(C)** Average paired pulse ratio for control (1.83 ± 0.15, n = 9) and Cre:GleFP+ neurons (1.70 ± 0.22, n = 9; t(16) = 0.4649, p = 0.6483, t test). Sample trace bars indicate 100ms, 30pA. **(D)** Averaged whole-cell LTP experiments and representative traces (50ms, 30pA). Summary of average percentage potentiation relative to baseline; control neurons (193.9 ± 22.1 %, n = 12), Cre:GFP+ neurons (196.6 ± 15.1 %, n = 12; t(22) = 0.09961, p = 0.9216, t test). Cumulative distribution of experiments.

### Single-neuron genetic deletion of serine racemase upregulates GluN2B in P45-70 CA1

We next sought to understand the discrepancy between the reduced NMDAR-EPSC amplitude and, in contrast to previous studies, the lack of effect on LTP (Basu et al., 2009; Benneyworth et al., 2012; Balu et al., 2013; Perez et al., 2017). We hypothesized that the reduced NMDAR-EPSC amplitude might be due to a decreased synaptic co-agonist concentration and thus less occupancy of the NMDAR co-agonist sites. To test this, we washed a saturating concentration of exogenous D-serine onto slices while recording from control and Cre+ neurons. If synaptic NMDARs had reduced co-agonist saturation in Cre+ relative to control cells, the D-serine wash would be predicted to cause a greater enhancement of the NMDAR-EPSCs in the Cre+ cells. However, we found that there was no significant difference in the potentiation of Cre+ and control cells in response to 100 μM D-serine (Fig 6A; control: 125.6 ± 4.1 %, n = 14; Cre+: 117.4 ± 14.4 %, n = 7; t(19) = 0.7147, p = 0.4835, t test).

**Figure 6:**
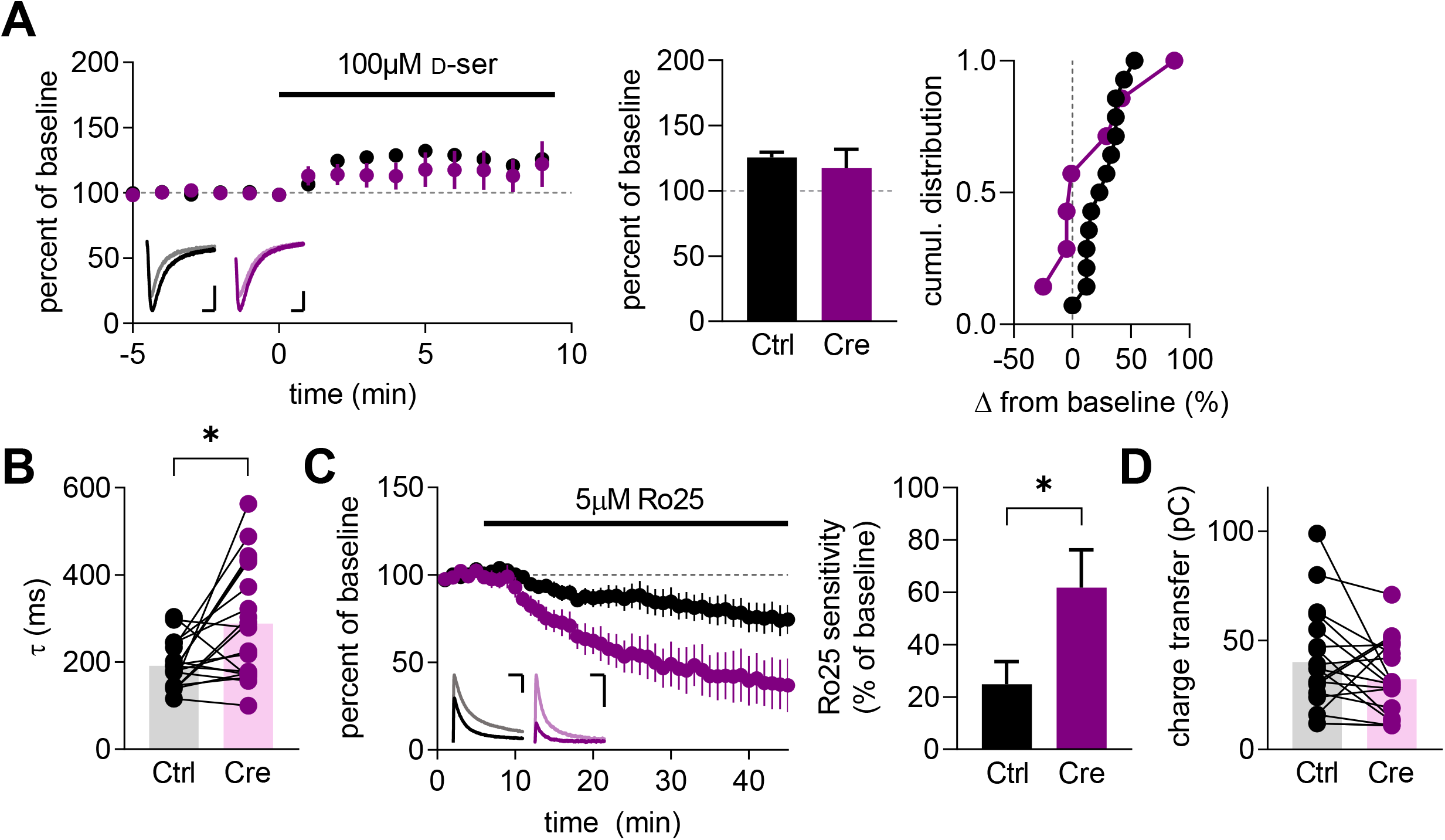
Single-neuron SR deletion increases synaptic GluN2B. **(A)** Averaged whole-cell D-serine wash experiments and representative traces (50ms, 30pA). Summary of average percentage potentiation relative to baseline; control neurons (125.6 ± 4.1 %, n = 14), Cre:GFP+ neurons (117.4 ± 14.4 %, n = 7; t(19) = 0.7147, p = 0.4835, t test). Cumulative distribution of experiments. **(B)** Average decay kinetics of NMDAR-EPSCs for control (191.1 ± 12.1 ms, n = 19) and Cre:GFP+ neurons (288.6 ± 30.6 ms, n = 19) from paired recordings in Fig 5A (t(18) = 3.040, p = 0.0070, paired t test). **(C)** Averaged whole-cell Ro25 wash experiments and representative traces (200ms, 50pA). Summary of average percentage of current sensitive to Ro25 wash; control neurons (24.8 ± 8.7 %, n = 6), Cre:GFP+ neurons (61.2 ± 14.5 %, n = 4; t(8) = 2.340, p = 0.0474, t test). **(D)** Average charge transfer of NMDAR-EPSCs for control (40.2 ± 5.4 pC, n = 19) and Cre:GFP+ neurons (32.3 ± 3.9 pC, n = 19) from paired recordings in Fig 2A (t(18) = 1.518, p = 0.1464, paired t test).

Interestingly, the decay kinetics of NMDAR-EPSCs were found to be significantly longer in Cre+ neurons (Fig 6B; control: 191.1 ± 12.1 ms, n = 19; Cre+: 288.6 ± 30.6 ms, n = 19; t(18) = 3.040, p = 0.0070, paired t test). Since prolonged decay of synaptic NMDAR-EPSCs likely indicates an increased proportion of the GluN2B subunit, we tested this pharmacologically with the GluN2B-selective inhibitor Ro25-6981 (Ro25) (Fischer et al., 1997). We found Cre+ neurons were significantly more sensitive than control neurons to 5 μM Ro25 (Fig 6C; control: 24.8 ± 8.7 %, n = 6; Cre+: 61.2 ± 14.5 %, n = 4; t(8) = 2.340, p = 0.0474, t test), demonstrating an increase in the synaptic GluN2B/GluN2A ratio.

GluN2B-containing NMDARs have lower peak open probability compared to GluN2A-containing NMDARs (Chen et al., 1999; Erreger et al., 2005; Gray et al., 2011). Though because their single channel conductances are identical (Stern et al., 1992), a reduction in the macroscopic EPSC amplitude could represent similar numbers of synaptic NMDARs. Indeed, the combination of reduced peak amplitude with the prolonged decay kinetics resulted in no significant change in charge transfer of Cre+ neurons (Fig 6D; control: 40.2 ± 5.4 pC, n = 19; Cre+: 32.3 ± 3.9 pC, n = 19; t(18) = 1.518, p = 0.1464, paired t test) consistent with a similar overall number of synaptic NMDARs. Overall, this increase in the GluN2B/GluN2A ratio may represent a homeostatic mechanism to maintain synaptic plasticity.

### Single-neuron genetic deletion of serine racemase impairs LTP in P23-39 CA1

Since the normal LTP in P45-70 mice might be due to homeostatic increases in GluN2B in the chronic absence of SR, we next examined an intermediate timepoint when D-serine is expected to be the primary synaptic co-agonist (Le Bail et al., 2015), but compensatory changes may not have yet occurred. Examining mice from the 4^th^-5^th^ week of life (P23-39), we found that the NMDAR-EPSCs were surprisingly increased in Cre+ neurons (Fig 7A; control: 91.7 ± 27.7 pA, n = 9; Cre+: 140.8 ± 26.3 pA, n = 9; t(8) = 2.789, p = 0.0236, paired t test). However, this EPSC increase was not accompanied by a significant change in decay time (Fig 7B; control: 200.8 ± 19.4 ms, n = 7; Cre+: 268.3 ± 60.4 ms, n = 7; t(6) = 1.510, p = 0.1818, paired t test). Similar to other timepoints, AMPAR-EPSCs (Fig 7D; 60.8 ± 12.7 pA, n =11; Cre+: 73.4 ± 10.4 pA, n = 11; t(10) = 0.9095, p = 0.3845, paired t test) and PPR (Fig 7E; control: 2.06 ± 0.21, n = 5; Cre+: 2.05 ± 0.12; t(8) = 0.01663, p = 0.9871, t test) were unchanged. Interestingly, LTP was completely eliminated in Cre+ neurons at P23-39 (Fig 7F; control: 140.2 ± 15.4 %, n = 6; Cre+: 94.7 ± 6.2 %, n = 6; t(10) = 0.0207, p = 0.0207, t test), demonstrating the neuronal SR cell-autonomously regulates NMDAR function and synaptic plasticity in CA1 pyramidal cells.

**Figure 7:**
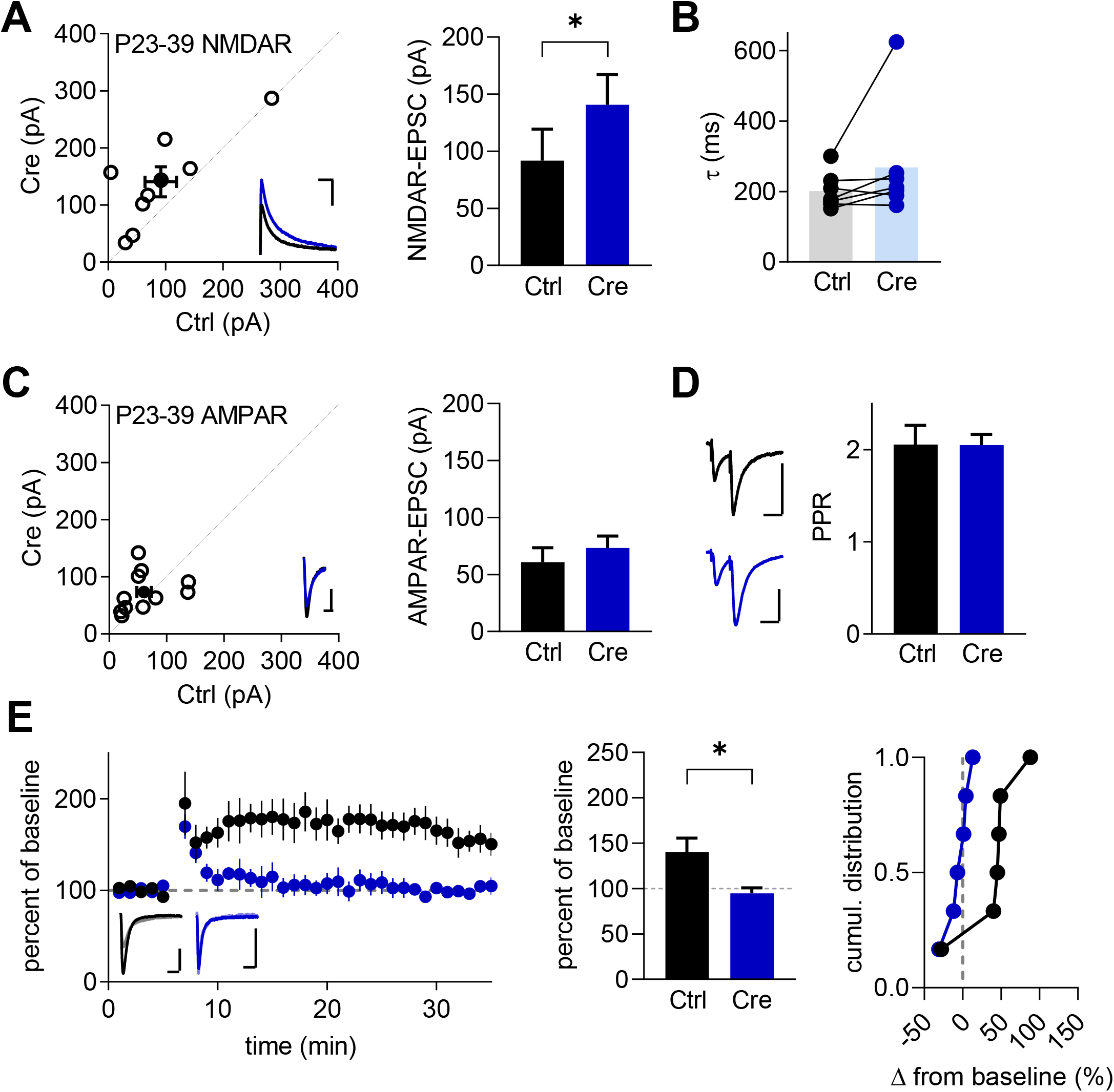
Loss of LTP at P23-39 following single-neuron SR deletion. **(A)** Scatterplot of paired neuronal recordings of P23-39 NMDAR-EPSCs (open circles) and averaged pair ± SEM (solid circle). Sample trace bars indicate 200ms, 30pA. Average NMDAR-EPSC amplitudes for control (91.7 ± 27.7 pA, n = 9) and Cre:GFP+ neurons (140.8 ± 26.3 pA; t(8) = 2.789, p = 0.0236, paired t test). **(B)** Average decay kinetics of NMDAR-EPSCs for control (200.8 ± 19.4 ms, n = 7) and Cre:GFP+ neurons (268.3 ± 60.4 ms, n = 7) from paired recordings in A (t(6) = 1.510, p = 0.1818, paired t test). **(C)** Scatterplot of paired neuronal recordings of P23-39 AMPAR-EPSCs (open circles) and averaged pair ± SEM (solid circle). Sample trace bars indicate 200ms, 30pA. Average AMPAR-EPSC amplitudes for control (60.8 ± 12.7 pA, n =11) and Cre:GFP+ neurons (73.4 ± 10.4 pA, n = 11; t(10) = 0.9095, p = 0.3845, paired t test). **(D)** Average paired pulse ratio for control (2.06 ± 0.21, n = 5) and Cre:GFP+ neurons (2.05 ± 0.12; t(8) = 0.01663, p = 0.9871, t test). Sample trace bars indicate 100ms, 30pA. **(E)** Averaged whole-cell LTP experiments and representative traces (50ms, 30pA). Summary of average percentage potentiation relative to baseline; control neurons (140.2 ± 15.4 %, n = 6), Cre:GFP+ neurons (94.7 ± 6.2 %, n = 6; t(10) = 0.0207, p = 0.0207, t test). Cumulative distribution of experiments.

## Discussion

Despite the recognized importance of D-serine in NMDAR function and synaptic plasticity, our understanding of the regulation of D-serine synthesis, release, and degradation remains quite limited. Indeed, even the cellular source of D-serine has been contested (Wolosker et al., 2016; Papouin et al., 2017; Wolosker et al., 2017). Early studies suggested that D-serine is exclusively synthesized and released by astrocytes (Schell et al., 1995; Schell et al., 1997; Wolosker et al., 1999) leading to the classification of D-serine as a gliotransmitter (Wolosker et al., 2002; Miller, 2004; Panatier et al., 2006). More recent studies, using the SR knockout mice as controls, have strongly supported a predominantly neuronal localization [recently reviewed by (Wolosker et al., 2016)]. In this study, we demonstrate for the first time a cell-autonomous role for neuronal SR in regulating synaptic NMDAR function. Specifically, we find that single-neuron genetic deletion of SR impairs LTP, although LTP is restored later with a concomitant upregulation of GluN2B. Furthermore, in agreement with previous studies in cultured neurons (Ma et al., 2014; Lin et al., 2016), we found that SR localizes to the apical dendrites and the post-synaptic density *in situ* in hippocampal CA1 pyramidal neurons. We have also identified D-serine in dendrites and postsynaptic compartments by immunogold EM. Together, these results provide strong evidence for the neuronal localization and cell-autonomous function of SR in the intact hippocampus. In addition, these findings together suggest a possible autocrine mode of D-serine action at synapses.

### Regulation of LTP by postsynaptic serine racemase

We demonstrate that single-neuron SR deletion cell-autonomously impairs LTP in P23-39 mice. Deficits in D-serine regulation have been repeatedly implicated in reduced synaptic LTP. Germline SR knockout mice show reduced LTP in CA1 (Basu et al., 2009; Balu et al., 2016; Neame et al., 2019), dentate gyrus (Balu et al., 2013), and lateral amygdala (Li et al., 2013). Furthermore, neuron-specific SR knockout mice also show reduced LTP in CA1 while astrocyte-specific SR knockout mice have normal LTP (Benneyworth et al., 2012; Perez et al., 2017). Similarly, acute enzymatic depletion of D-serine reduces the magnitude of LTP in CA1 (Yang et al., 2003; Papouin et al., 2012; Rosenberg et al., 2013; Le Bail et al., 2015), visual cortex (Meunier et al., 2016), nucleus accumbens (Curcio et al., 2013), and lateral amygdala (Li et al., 2013). Together, these studies suggest that neuronally-derived D-serine is crucial for synaptic plasticity, and the cell-autonomous nature of the LTP loss seen here suggests a possible autocrine mode of D-serine action (discussed below).

Interestingly, we found that LTP was restored in P45-P70 mice, possibly through a homeostatic process. This restoration of LTP was associated with a significant upregulation of synaptic GluN2B. Indeed, a prolongation of the NMDAR-EPSC decay kinetics, consistent with an upregulation of synaptic GluN2B, has been previously reported in the germline SR knockout mice (Basu et al., 2009), and GluN2B is known to promote LTP through its unique array of c-tail interacting proteins (Foster et al., 2010). Importantly, the enhancement of GluN2B subunits in CA1 could directly compensate for a lack of D-serine. GluN2A and GluN2B allosterically regulate co-agonist potency at the GluN1 glycine binding site (Priestley et al., 1995; Madry et al., 2007; Chen et al., 2008; Maolanon et al., 2017) with a two-to five-fold higher potency of co-agonists at GluN2B-containing NMDARs. Therefore, enhancement of GluN2B could compensate for a loss of D-serine given a smaller but stable pool of synaptic glycine. Indeed, the increased affinity of GluN2B-containing NMDARs for co-agonist could explain the lack of NMDAR saturation changes in the P45-P70 mice. Other groups using the SR germline KO or the broad neuronal SR deletion, however, did not observe this restoration of LTP (Basu et al., 2009; Balu et al., 2016; Perez et al., 2017; Neame et al., 2019). These studies examined mice from P21 to 5 months old, thus it is unlikely that our results are simply due to a previously uncharacterized age, but could be a result of the single-neuron approach. Perhaps there is some degree of D-serine spillover from neighboring neurons that is sufficient to fully activate the higher-affinity GluN2B-containing NMDARs and restore LTP, whereas when SR is deleted from all neurons, this spillover is eliminated.

Alternatively, the increase in synaptic GluN2B might solely be an associated finding. For example, NMDAR synaptic stability may be regulated by co-agonist composition in a subunit-specific manner. Co-agonist binding primes NMDARs for internalization and recycling (Nong et al., 2003), and D-serine administration onto dissociated cortical cultures increases the rate of GluN2B surface diffusion and decreases residence at postsynaptic sites (Papouin et al., 2012; Ferreira et al., 2017). Thus, loss of D-serine at synaptic sites by the removal of SR may alter the balance of synaptic GluN2 subunits through changes in trafficking mechanisms. Indeed, the surprising increase in NMDAR-EPSCs at P23-39, even with the absence of LTP, might be indicative of early alterations of subunit trafficking.

### Autocrine mode of D-serine action?

The postsynaptic localization of SR and the cell-autonomous regulation of NMDAR function by neuronal SR suggests the local postsynaptic release and autocrine mode of D-serine action at synapses. For example, in addition to pyramidal neurons, SR and D-serine also localize to GABAergic neurons (Miya et al., 2008; Curcio et al., 2013; Balu et al., 2014; Lin et al., 2016; Takagi et al., 2020), arguing against D-serine being release presynaptically as a co-transmitter. Furthermore, recent work corroborates an autocrine mode of D-serine action via AMPAR and NMDAR activity-dependent regulation of SR at synapses. *In vitro*, SR forms a ternary complex with PSD95 and the AMPAR accessory protein stargazin and this interaction increases the membrane localization of SR and is associated with a decrease in SR activity (Ma et al., 2014). After activation of AMPARs, SR dissociates from this complex and drives D-serine synthesis (Ma et al., 2014). In culture, exogenous D-serine enhances association of SR with PSD-95 and GluN1 in co-immunoprecipitation experiments and this association is blocked by the NMDAR glycine-site antagonist 7-chlorokyneurenic acid (Lin et al., 2016). In addition, NMDAR activity leads to the SR palmitoylation increasing its membrane association and decreased its activity (Balan et al., 2009). These studies suggest tightly controlled regulation of postsynaptic SR activity. However, the physiological importance of these protein-protein interactions and posttranslational modifications in intact neuronal circuits is not known.

An autocrine mode of D-serine action is further supported by identification of D-serine transporters in neurons. Alanine-serine-cysteine transporter 1 (Asc-1) is a neutral amino acid transporter located in neurons (Helboe et al., 2003; Matsuo et al., 2004) that can mediate the bidirectional transport of D-serine while swapping with other small neutral amino acids (Fukasawa et al., 2000). Neuronal D-serine release via Asc-1 from cultured neurons by the addition of D-isoleucine stimulates the antiporter activity of Asc-1 (Rosenberg et al., 2013; Sason et al., 2017). Perfusion of 1 mM D-isoleucine onto acute hippocampal slices increased NMDAR-fEPSPs and enhanced LTP (Rosenberg et al., 2013) and inhibition of Asc-1 decreases both D-serine uptake and release and also inhibits LTP in CA1 (Sason et al., 2017). However, Asc-1 also transports other amino acids, including glycine and L-serine, which complicates interpretation of its effects (Fukasawa et al., 2000). Furthermore, the subcellular localization of Asc-1 is not known.

### High levels of non-synaptic serine racemase

As we show here (Fig 1 and 2), SR immunoreactivity is curiously high in non-synaptic locations, notably the soma and dendrites, but for what purpose? Evidence suggests that both nuclear and dendritic localization of SR downregulates its activity. For example, following apoptotic insult, SR translocates to the nucleus independent of NMDAR activity where its racemase activity is inhibited to limit apoptotic damage (Kolodney et al., 2015). Interestingly, in addition to its function as a racemase, SR can also function as an eliminase, catalyzing the α,β-elimination of water from both L-serine (Strisovsky et al., 2003) and D-serine (Foltyn et al., 2005) to form pyruvate. At least *in vitro*, SR produces 3-fold more pyruvate than D-serine, suggesting that the eliminase activity is dominant (Panizzutti et al., 2001; Foltyn et al., 2005). Indeed, these dueling activities of SR may function to limit intracellular D-serine levels (Foltyn et al., 2005). Because its racemase activity can be controlled by post-translational modifications and protein-protein interactions (Balan et al., 2009; Foltyn et al., 2010; Ma et al., 2014), and racemase and eliminase activity are differentially controlled by coenzyme availability (Strisovsky et al., 2003), SR may have pleotropic roles dependent on subcellular localization. Nevertheless, this study does not ultimately distinguish between SR racemization and elimination.

In summary, our data show the postsynaptic localization of SR in hippocampal CA1 pyramidal neurons and the cell-autonomous regulation of NMDARs by neuronal SR. These results support an autocrine mode of D-serine following postsynaptic release. Indeed, the concept of postsynaptic release of neuromodulators is not new. For example, brain-derived neurotrophic factor (BDNF) is released postsynaptically during synaptic plasticity (Harward et al., 2016; Hedrick et al., 2016). However, further studies are needed to identify the mechanisms regulating D-serine postsynaptic release and termination of D-serine action within the synaptic cleft.

## Acknowledgements

This work was supported by: Jeane B. Kempner Postdoctoral Fellowship Award (OF); Whitehall Foundation #2018-05-107 (DTB); T32MH082174 (EB); BrightFocus Foundation A2019034S (DTB); R03AG063201 (DTB); a subcontract of R01NS098740 (DTB); R01MH118497 (MLM); R01MH05190 (JTC); R21MH116315 (JAG); and R01MH117130 (JAG). We would like to thank Haley Martin, Zaiyang “Sunny” Zhang, and Casey Sawyer for their assistance in mouse breeding and genotyping. Human tissue was obtained from the NIH NeuroBioBank.

